# p-IgGen: A Paired Antibody Generative Language Model

**DOI:** 10.1101/2024.08.06.606780

**Authors:** Oliver M. Turnbull, Dino Oglic, Rebecca Croasdale-Wood, Charlotte M. Deane

## Abstract

A key challenge in antibody drug discovery is designing novel sequences that are free from developability issues - such as aggregation, polyspecificity, poor expression, or low solubility. Here, we present p-IgGen, a protein language model for paired heavy-light chain antibody generation. The model generates diverse, antibody-like sequences with pairing properties found in natural antibodies. We also create a finetuned version of p-IgGen that biases the model to generate antibodies with 3D biophysical properties that fall within distributions seen in clinical-stage therapeutic antibodies.

## Introduction

Antibodies play a crucial role in the immune response and are an increasingly important class of therapeutic (18). They consist of two sets of heavy and light chains with antigen binding mediated by the Fv region of each chain (VH and VL respectively) (3). The majority of the diversity in antibodies is located in six hypervariable loops within the Fv region known as complementarity-determining regions (CDRs). The light chain and heavy chain each contain 3 CDR loops (CDRL 1-3 and CDRH 1-3).

Modern antibody drug discovery typically relies on large libraries of paired variable heavy (VH) and variable light (VL) sequences which are screened for affinity against a target (23). However, such libraries often contain sequences with developability issues (10) (i.e. propensity for aggregation, polyspecificity, poor expression, or low solubility) which can result in potential therapeutics being discarded or requiring engineering later in the pipeline (18). These developability properties depend on the combination of the heavy and light chain.

Protein language models (PLMs) have proven effective in a variety of tasks. Masked PLMs are trained with the task of predicting masked-out tokens (in the case of PLMs, residues) in a sequence and thereby learn a rich representation of protein sequences. Masked PLMs have proven effective for tasks such as property prediction (14) and suggesting evolutionarily plausible mutations (8). Auto-regressive (AR) models are trained to predict the next residue in the sequence and during inference are able to generate full sequences. AR PLMs have demonstrated the ability to generate novel and diverse protein sequences (7, 15, 21).

Here, we present paired-IgGen (p-IgGen), an auto-regressive PLM trained on both unpaired and paired antibody sequences. p-IgGen generates diverse and antibody-like paired sequences, as measured with a variety of sequence and structure-based metrics. Our extensive validation indicates the paired sequences generated by p-IgGen follow similar pairing patterns to those seen in natural paired sequences. To make best use of available antibody sequence data we use a pretraining regime capable of ingesting the large corpus of unpaired sequences (∼250M) followed by finetuning on the smaller but more biologically relevant paired sequence data (∼1.8M). The model can also be biased to generate sequences with desired properties through finetuning. Here, we bias the model to generate antibodies with 3D biophysical properties that fall within distributions seen in clinical-stage therapeutic antibodies, as predicted by the structure-based developability predictor the Therapeutic Antibody Profiler (TAP) (17), which we release as ‘developable p-IgGen’. Finally, we show that p-IgGen outperforms other antibody language models on zero-shot prediction benchmarks, demonstrating robust sequence representations.

## Results

### Paired Antibody Language Model

p-IgGen is an autoregressive decoder-only language model using a GPT -2-like architecture (2), see Methods for full architecture and training details. We trained p-IgGen in a two-step procedure, first pretraining on the much larger available dataset of unpaired sequences. ‘IgGen’ was trained on a filtered set of 117M VL and 140M VH sequences taken from the Observed Antibody Space (OAS) (16), see Methods for full filtering and tokenisation details. ‘p-IgGen’ was then trained by finetuning IgGen on a set of 1.8M paired VH/VL sequences taken from OAS.

### p-IgGen generates novel, realistic and diverse paired sequences

We evaluated the sequences generated by p-IgGen using a comprehensive set of in silico metrics, demonstrating that its generated sequences are unique, diverse, and antibody-like. By comparing these metrics against a test set of natural paired sequences we establish that the distributions of generated sequences are very similar to those of natural sequences.

We found that sequences generated by p-IgGen were as similar to natural sequences as natural sequences are to each other, as measured by Hamming distance (Table 1). Generated sequences also show a similar sequence identity to both training and validation sets, indicating that the model has not overfit to the training data (see Supplementary Information). Following Shin et al. (20) we examined the diversity of generated sequences using the cosine similarity with each sequence’s nearest neighbour. At a sampling temperature of 1.2, the generated sequences were as diverse as natural sequences (see Table 1). This diversity can also be tuned by adjusting the sampling temperature.

**Table 1.** Comparison of Metrics between p-IgGen Sequences, Randomly Paired p-IgGen Sequences, and Natural Sequences. We compared a variety of metrics investigating the novelty, diversity, and validity of sequences. As a control, we randomly paired the VH and VL chains from the ‘p-IgGen Sequences’ set. Natural sequences were taken from the training set. Diversity is measured by the mean cosine distance between sequences. VH-VL mutation rate correlation was calculated by looking at the correlation between the number of mutations away from germline in the VH and VL chains.”

| Metric | p-IgGen Sequences | Randomly Paired p-IgGen Sequences | Natural Sequences |
| --- | --- | --- | --- |
| Mean VH Hamming Distance to Val Set | 11.3 | 11.3 | 11.3 |
| Mean VL Hamming Distance to Val Set | 10.3 | 10.3 | 9.5 |
| Diversity | 0.16 | 0.17 | 0.16 |
| ESM-2 Likelihood | -0.33 | -0.33 | -0.33 |
| VH-VL Mutation Rate Correlation | 0.52 | -0.04 | 0.51 |

Generated sequences show a similar distribution of ESM-2 (12) likelihoods, suggesting that they are just as ‘protein-like’ as natural sequences. To assess if they were antibody-like, we aligned and numbered all sequences with the antibody-numbering tool ANARCI (6) which successfully identified a heavy and light chain for all sequences. The distribution of CDR lengths was also examined and found to be closely aligned with natural sequences (see Supplementary Information). We also tested whether the sequences could be structurally modelled using ABodyBuilder2 (ABB2) (1) and found similar confidence values as those seen for natural sequences (see Supplementary Information).

Finally, we investigated whether the generated sequences show VH/VL pairing characteristics similar to those observed in natural sequences. Naturally paired sequences show a correlation in the mutation rates of the heavy and light chains, relative to their respective germline sequences. We found that generated sequences from p-IgGen also display a similar correlation, while no such correlation is observed when the same generated sequences are randomly paired together (see Table 1). This suggests that sequences generated by pIgGen are not only antibody-like but also have biologically plausible VH/VL pairings.

The validation described above of p-IgGen’s generated sequences spans a broad array of tests: assessing sequence identity and diversity, protein-likeness, antibody-specific properties (including ANARCI numbering and CDR length distributions), and conducting structural modelling with ABB2. Together, these tests confirm that p-IgGen generates novel and diverse sequences that appear just as antibody-like as natural sequences. This comprehensive validation confirms p-IgGen’s potential to generate novel, realistic, and diverse paired antibody libraries.

### Generation can be biased towards antibodies with desired sequence and structure-based properties

Having verified that p-IgGen can create diverse, realistic, and previously unseen antibodies, we then investigated whether the generation space could be restricted to antibodies with desirable developability properties. The approach we took was to fine-tune p-IgGen on a set of antibodies with the desired properties. This has the advantage of being very simple to implement; it does not require a differentiable property predictor or reinforcement learning. As a case study, we fine-tuned on a set of developable antibodies, as predicted by the Therapeutic Antibody Profiler (TAP) tool (17). Specifically, we structurally modelled all 1.8M paired sequences using ABB2 and ran these structures through TAP. We defined an antibody as ‘developable’ if it had all green flags for the four structure-based TAP metrics (PSH, PPC, PNC, and SFvCSP) (see Methods for full details). We used this ‘safe’ set of 909,790 sequences to fine-tune paired p-IgGen to create a “developable p-IgGen”. The developable p-IgGen is therefore trained in a three-step process - first pretrained on unpaired data, then finetuned on paired sequences, and finally finetuned on highly developable sequences.

Finetuning on the developable set shifted the distribution of the 3D biophysical properties of generated antibodies despite being trained on sequence alone. We saw a significant reduction in the proportion of amber and red-flagged antibodies for all metrics for the developable model relative to the paired model (Figure 1). The diversity of generated antibodies was still maintained, as measured by intraset diversity and sequence identity (see Supplementary Information).

**Fig. 1.**
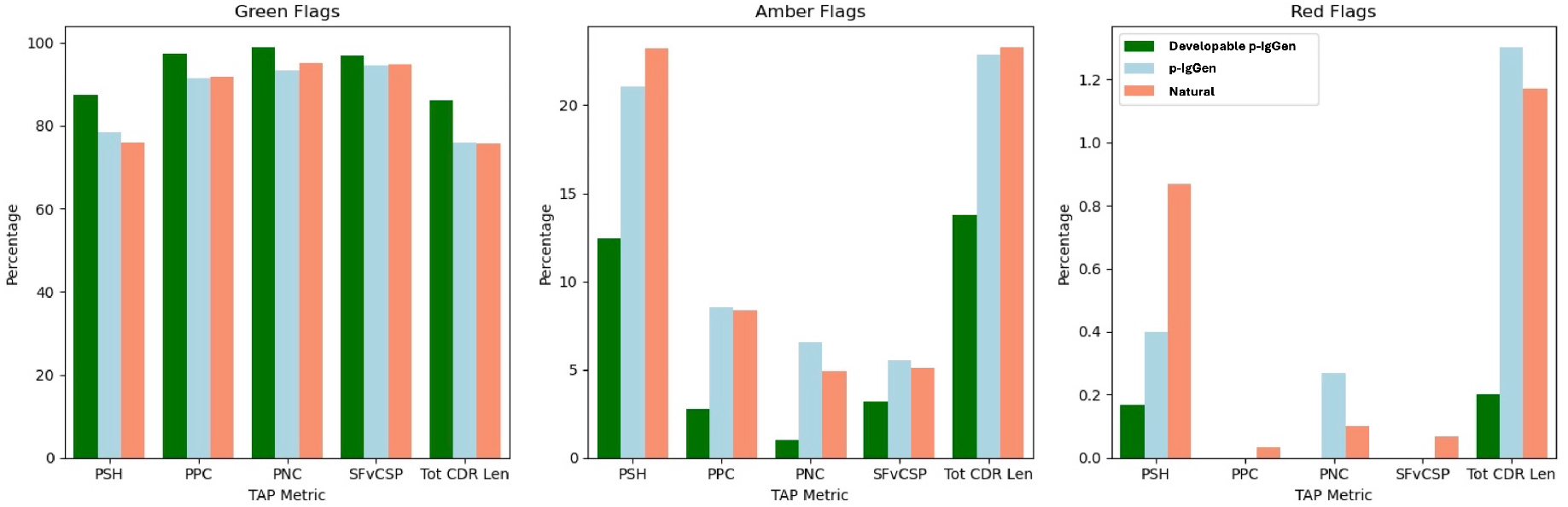
Developable p-IgGen shows favourable TAP flagging compared to p-IgGen on both structure-based and sequence-based metrics. We generated 3000 sequences from p-IgGen, and developable p-IgGen and sampled 3000 sequences from the paired OAS validation set. All of these sequences were structurally modelled with ABB2 and were then run through TAP to generate developability flags for the four structure-based metrics (PSH, PPC, PNC, SFvCSP) and the total CDR length. Green flagged sequences represent low developability risk, amber flagged are medium risk, and red flagged are high risk.

### p-IgGen achieves state-of-the-art performance of 0-shot tasks

Finally, to understand whether our models were learning meaningful representations of antibodies, we tested their ‘zero-shot’ prediction performance on two antibody fitness datasets. We used a deep mutational scan dataset of 4275 anti-VEGF antibodies (11) to assess zero-shot prediction of expression, and a curated dataset of antidrug antibody responses for 217 therapeutic antibodies for immunogenicity prediction (13). For zero-shot accuracy, we looked at the Pearson correlation of the perplexity of sequences under the given model with the fitness metric being assessed.

For model testing and comparison, we used the Fitness Landscapes for Antibodies (FLAb) testing suite (4). FLAb offers benchmark results for various state-of-the-art models, both sequence-based (IgLM (21), AntiBERTy (19), ProGen (15)) and structure-based (ProteinMPNN (5), ESM-IF (9)). However, FLAb lacks benchmarks for structure-based models on the expression dataset. For both datasets, we found that p-IgGen outperformed IgGen, while p-IgGen and developable p-IgGen performed similarly. For the expression dataset, p-IgGen outperformed all other antibody-specific LMs (AntiBerty, IgLM, and ProGen OAS) (see Supplementary Information). The ProGen general protein LMs outperformed p-IgGen for expression prediction, but even the smallest ProGen model has more than 7.5X the number of parameters of p-IgGen and requires significantly more compute to train. The superior performance of general protein language models for expression suggests the evolutionary patterns learnt during training on diverse proteins may be important for the more general problem of protein expression. For the immunogenicity dataset, p-IgGen outperformed all other methods, with the same performance for both the developable p-IgGen model and the p-IgGen model (Table 2).

**Table 2.**
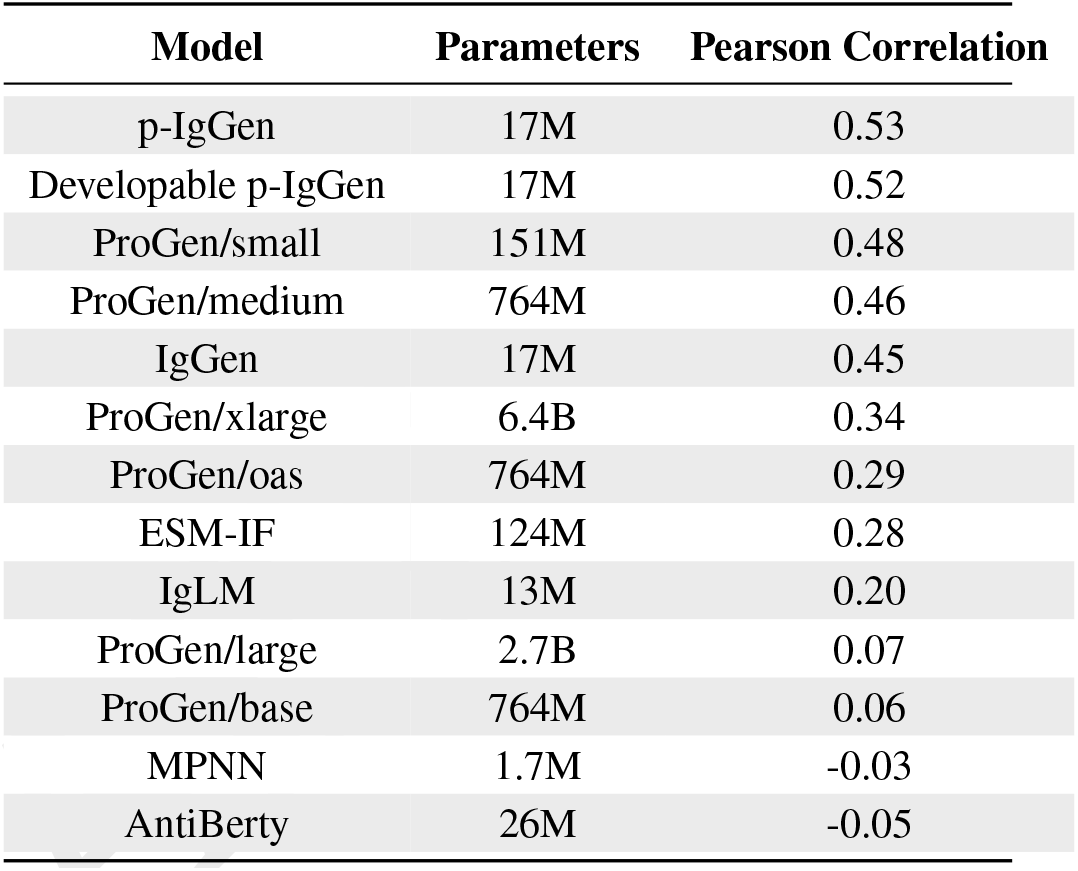
p-IgGen and developable p-IgGen outperform all other models for zero-shot prediction of immunogenicity. Language models and inverse folding models (ESM-IF and MPNN) were evaluated for zero-shot prediction of immunogenicity using a dataset of antidrug antibody responses for 217 therapeutic antibodies curated by Marks et al. (13) using FLAb. Results are ordered by Pearson’s correlation (best to worst).

| Model | Parameters | Pearson Correlation |
| --- | --- | --- |
| p-IgGen | 17M | 0.53 |
| Developable p-IgGen | 17M | 0.52 |
| ProGen/small | 151M | 0.48 |
| ProGen/medium | 764M | 0.46 |
| IgGen | 17M | 0.45 |
| ProGen/xlarge | 6.4B | 0.34 |
| ProGen/oas | 764M | 0.29 |
| ESM-IF | 124M | 0.28 |
| IgLM | 13M | 0.20 |
| ProGen/large | 2.7B | 0.07 |
| ProGen/base | 764M | 0.06 |
| MPNN | 1.7M | -0.03 |
| AntiBerty | 26M | -0.05 |

## Conclusions

In this work, we have presented and extensively validated p-IgGen, an antibody language model capable of producing realistic paired sequences and achieving state-of-the-art performance on zero-shot tasks. The ability to finetune p-IgGen to produce sequences with desired biophysical properties while preserving diversity highlights its applicability to high throughput antibody drug discovery.

## Methods

### Model and Training

We trained three models: IgGen, p-IgGen, and developable p-IgGen. IgGen was pretrained on unpaired sequences and finetuned on paired sequences to give p-IgGen. p-IgGen was further finetuned on a set of developable sequences to give developable p-IgGen. All models use the same autoregressive decoder-only architecture based on GPT-2 (2) with the addition of rotary positional embedding (22), implemented in PyTorch. We used 3 attention layers, each with 12 attention heads and an embedding size of 768, the feed-forward layers had a dimension of 2048, for a total of 17,349,888 parameters. Sequences were tokenised at the residue level, with a special token added to the start (“1”) and end (“2”) of each sequence. For the paired models, the light chain was concatenated to the heavy chain. During training, we randomly provided each sequence either in the forward or reverse direction. By training in both the ‘forward’ and ‘reverse direction, a heavy chain can be generated given a light chain and vice versa during inference.

All training was performed using the Adam optimiser with a cosine learning rate scheduler. IgGen was trained for 20 epochs on 5 A100 GPUs with a learning rate of 1E-4, a local batch size of 512 and 4 gradient accumulation steps. p-IgGen was trained by finetuning all layers of IgGen for 3 epochs using a batch size of 256 and a learning rate of 1E-5 on an A100 GPU. 3 epochs was chosen as the model showed an understanding of paired sequences while showing less forgetting of unpaired sequences in comparison to further trained models. Finally, developable p-IgGen was trained by finetuning all layers of p-IgGen for 2 epochs with the same hyperparameters as used to train p-IgGen.

### Model Sampling

We found that using a sampling temperature of 1.2, a top-p of 0.95, and then discarding the bottom 5% of generated sequences according to the model likelihood best represented natural antibody sequences in terms of mutation rate and diversity while remaining valid. For developable p-IgGen, we increased the sampling temperature to 1.25 to maintain diversity. We used 2000 sequences for all comparisons.

### Data

Models were trained using antibody sequences taken from the Observed Antibody Space (OAS) (16). Only human sequences were used, and sequences were filtered to reduce redundancy and to remove sequences which likely contained PCR sequencing errors (see Supplementary Information for full filtering details). For the unpaired dataset, this resulted in 117,431,915 VL and 130,246,252 VH sequences. The filtered paired dataset consists of 1,800,545 VH/VL sequences.

For the developable dataset, we structurally modelled all sequences from paired OAS using ABodyBuilder2 (1). The structures were then flagged for developability using TAP (17), which calculates five metrics which have been associated with poor developability. CDR sequences that had green flags for the four structure-based metrics (PSH, SFvCSP, PPC, and PNC) were classified as developable and included in the finetuning set. We did not filter on the CDR length metric, as this is sequence-based so generated sequences could be quickly and easily filtered for this. As p-IgGen had already been trained on the paired dataset, we ensured that we kept the same train / validation / test splits as used for training the paired model.

Datasets for zero-shot prediction tasks were taken from the FLAb repository (4). The immunogenicity set consists of the anti-drug antibody (ADA) response against 217 therapeutics curated by Marks et al. (13). The expression dataset is taken from Koenig et al. (11) and consists of a deep mutational scan of an anti-VEGF antibody, with 4275 sequences. There was no overlap between the paired zero-shot test sequences and the unpaired or paired training set sequences.

## Data Availability

The data underlying this article are available in the Observed Antibody Space (OAS), at https://opig.stats.ox.ac.uk/webapps/oas.

