## Appendix for "p-IgGen: A Paired Antibody Generative Language Model"

#### 1 Tokenisation Scheme

| Forward Tokenisation | Reverse Tokenisation |
|---|---|
| 1{VH}2 | 2{Reverse VH}1 |
| 1{VL}2 | 2{Reverse VL}2 |
| 1{VH}{VL}2 | 2{Reverse VL}{Reverse VH}1 |

Table 1: **Tokenisation Scheme** IgGen is provided with VH and VL sequences separately, while p-IgGen and developable p-IgGen are provided with the VL concatenated to the VH. All sequences are provided in the forward direction as well as reversed. During training, models are shown all sequences in both the forward and reverse direction.

#### 2 Dataset Filtering

For unpaired OAS, heavy and light sequences were filtered separately to remove identical sequences and any sequences marked by ANARCI [Dunbar and Deane, 2015] as having shorter than IMGT defined framework region 1 or 4, missing conserved cysteines, or containing unknown residues. The sequences were then further filtered for redundancy by clustering at 95% identity using linclust [Steinegger and Söding, 2018] with coverage mode 1 (target coverage). Within each cluster, we further clustered by identical CDRs and kept a random sample for each sub-cluster. We numbered sequences with the IMGT scheme using ANARCI [Dunbar and Deane, 2015] and used IMGT CDR definitions [Lefranc et al., 2003]. 117,431,915 VL and 130,246,252 VH sequences were used for further steps.

Paired OAS was filtered to remove sequences with missing conserved cysteine residues or with unknown residues. Sequences with deletions in framework regions were completed using AbLang [Olsen et al., 2022b]. This was not performed for unpaired sequences as a large amount of data was already available. Due to the smaller size of paired OAS, and the increased diversity relative to unpaired OAS due to the combination of both VH and VL chains for each sequence, we did not filter the sequences for redundancy, apart from ensuring no identical full VH/VL sequences were present. For the train, validation, and test splits, we clustered length matched CDRs at 95% sequence identity using cd-hit [Li and Godzik, 2006].

### 3 Sequence Validation

We generated samples using top-p sampling, as implemented in the HuggingFace Transformers library [Wolf et al., 2020], with a top-p value of 0.95, and a temperature value of 1.2 unless otherwise stated. We then calculated the model likelihood of generated sequences and discarded the bottom 5% of sequences. We numbered sequences using ANARCI [Dunbar and Deane, 2015] and the IMGT scheme [Lefranc et al., 2003] to identify the heavy and light chains and allow for other downstream analyses.

To assess the intraset diversity of the generated and test sequences, we calculated the pairwise cosine diversity of 3-mer subsequences of the paired sequences within each set, with the light chain concatenated after the heavy chain, using the scikit-learn library [Pedregosa et al., 2011]. We calculated the pseudo-log-likelihood of sequences using ESM2 [Lin et al., 2023], with the light chain concatenated after the heavy chain using the esm2_t12_35M_UR50D model hosted on HuggingFace [Wolf et al., 2020].

We used ANARCI-derived numbering and IMGT definitions to calculate the length distribution of the CDRs within the generated and natural sets. ANARCI annotations were also used for germline gene usage and sequence identity to germline sequences. We modelled all generated sequences using ABodyBuilder2 (ABB2) [Abanades et al., 2023] and extracted error estimates from the generated pdb files. For developability prediction, we ran the generated structures through the Therapeutic Antibody Profiler (TAP) [Raybould et al., 2019].

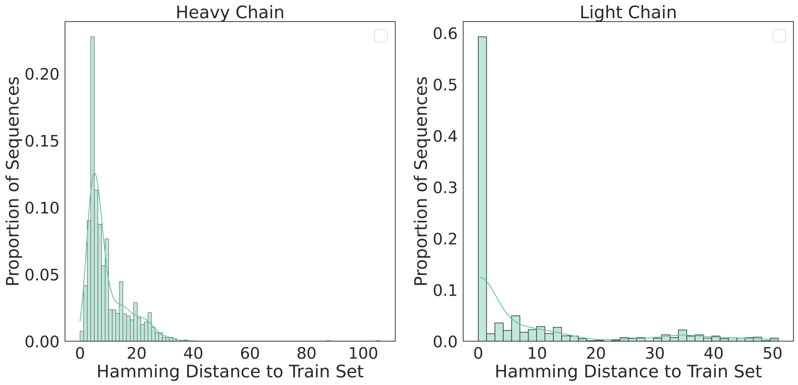


Figure 1: **Generated sequences do not show signs of overfitting to the training sequences.** We calculated the minimum Hamming distance of VH and VL from 2,000 sequences generated by p-IgGen with the paired OAS training set. VH and VL regions were extracted from the generated sequences using ANARCI. KDE lines show the smoothed distribution of the sequence identity data.

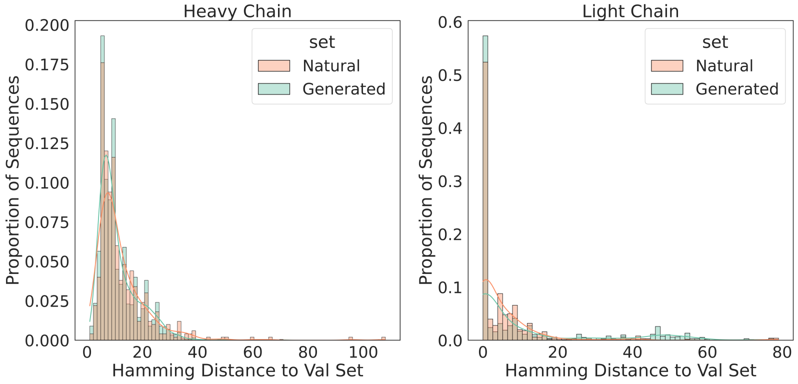


Figure 2: **Generated sequences show similar distances to validation set sequence as training set sequences do to validation set sequences.** We calculated the Hamming distance of VH and VL from 2,000 sequences generated by p-IgGen with the paired OAS validation set (”Generated”). We also calculated the sequence identity of a random sample of 2,000 OAS paired training set sequences to validation set sequences (”Natural”). VH and VL regions were extracted from the generated sequences using ANARCI. KDE lines show the smoothed distribution of the sequence identity data.

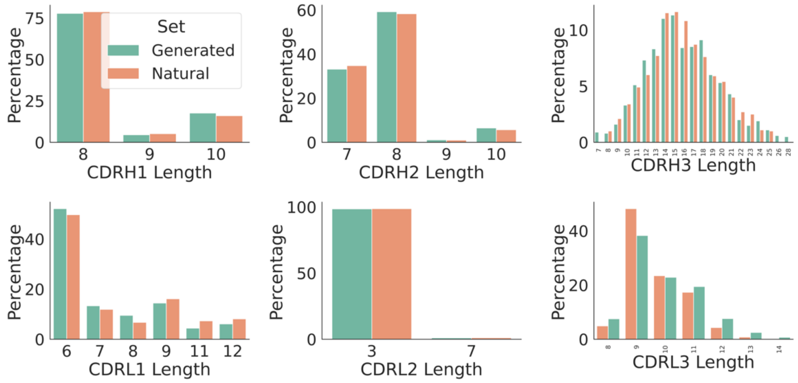


Figure 3: **Generated sequences show a similar distribution of CDR lengths to natural sequences.** We looked at the distribution of CDR lengths of sequences generated by p-IgGen ("Generated") compared to test set sequences from paired OAS ("Natural"). Lengths were determined using IMGT-defined CDR positions with IMGT numbering using ANARCI.

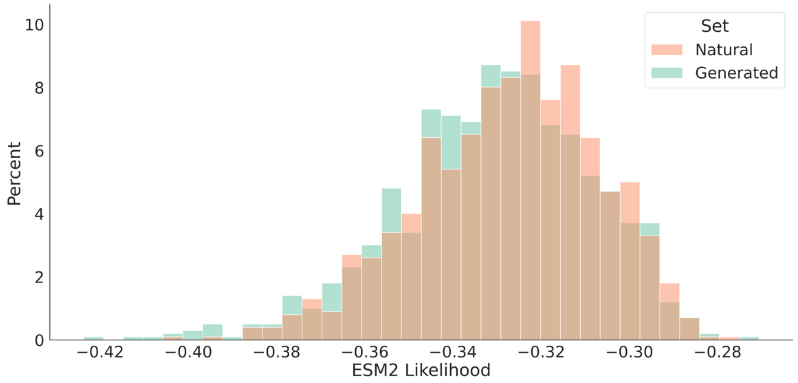


Figure 4: **Generated sequences have a similar distribution of ESM-2 log-likelihoods as natural sequences.** We calculated the log-likelihood of 2,000 full VH/VL sequences generated by p-IgGen ("Generated") as well as 2,000 sequences taken from the test set of paired OAS using the masked protein language model ESM-2.

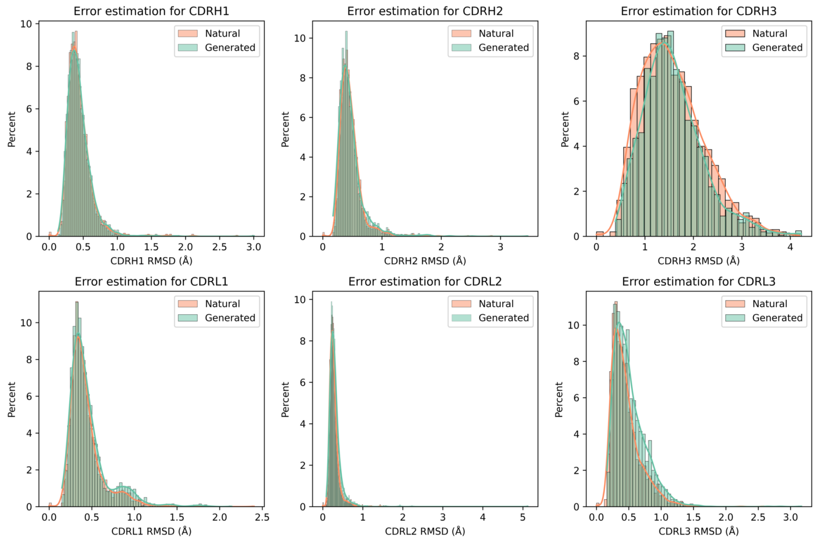


Figure 5: **Generated sequences have similar structural modelling error estimates as natural sequences.** We structurally modelled 2,000 generated and 2,000 natural sequences using ABB2. Per loop error estimates were produced by taking the mean ABB2 RMSD error estimate across residues in IMGT-defined CDR regions, as numbered by ANARCI.

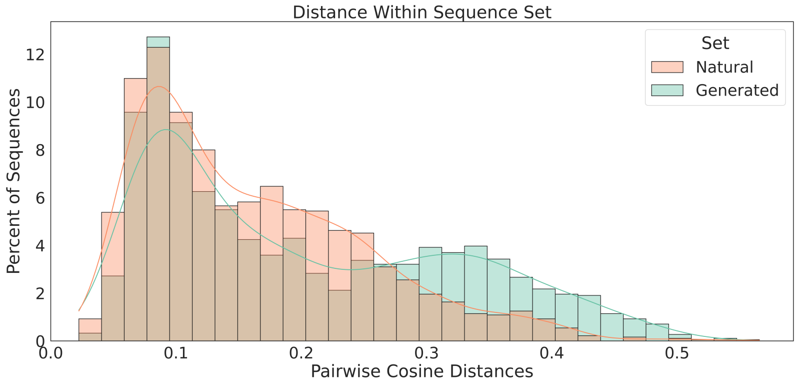


Figure 6: **Intraset diversity, measured by cosine distance, is similar for generated and natural sequences.** We calculated the highest pairwise cosine distance for 2,000 sequences generated from p-IgGen using a sampling temperature of 1.2 ("Generated"). This was compared to the highest pairwise cosine distance for 2,000 sequences sampled from the validation set of paired OAS ("Natural"). KDE lines show the smoothed distribution of the diversity data.

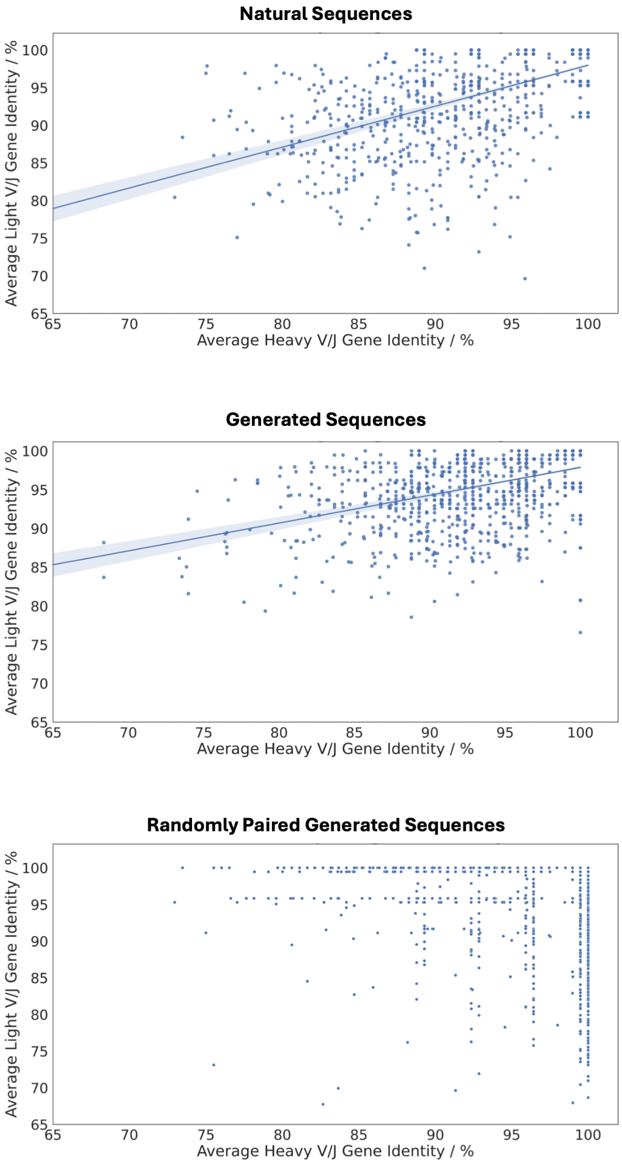


Figure 7: **Both natural sequences and paired sequences generated by p-IgGen show a correlation between the mutation rates of the VH and VL chains.** Average V/J gene identity to germline was used as a measure of mutation of the VH and VL chains, as reported by ANARCI. Natural sequences (taken from the paired OAS validation set) and sequences generated by p-IgGen ("Generated Sequences") show a strong correlation between the VH and VL mutation rates. No correlation is seen for generated sequences with randomly paired VH and VL chains.

### 4 Property Biasing

We found that after fine-tuning, a slightly higher sampling temperature of 1.25 was needed to achieve similar diversity to natural sequences. However, a lower diversity is also expected as we've restricted the generation space to developable antibodies.

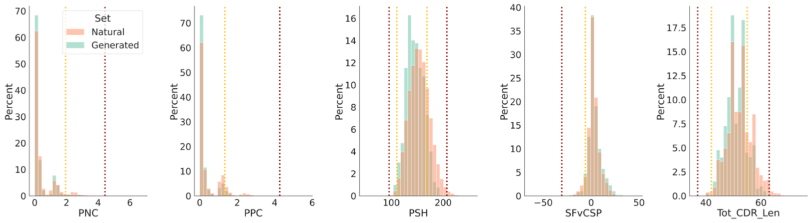


Figure 8: **Antibodies generated by developable p-IgGen show a favourable shift in the distribution of TAP metrics relative to natural sequences.** We generated and structurally modelled 2,000 sequences from developable p-IgGen using ABB2. We then ran TAP on the structural models to calculate the four structure-based metrics (PNC, PPC, PSH, and SFvCSP) and the total CDR length ("Generated"). We also calculated the TAP metrics for all paired OAS test set sequences ("Natural") using the same methodology.

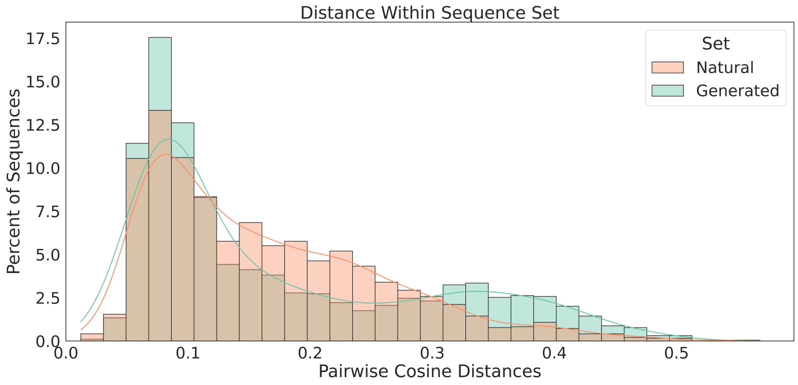


Figure 9: **Sequences generated by developable p-IgGen maintain their diversity.** We calculated the highest pairwise cosine distance for 2,000 sequences generated from developable p-IgGen using a sampling temperature of 1.25 ("Generated"). This was compared to the highest pairwise cosine distance for 2,000 sequences sampled from the validation set of developable paired OAS ("Natural").

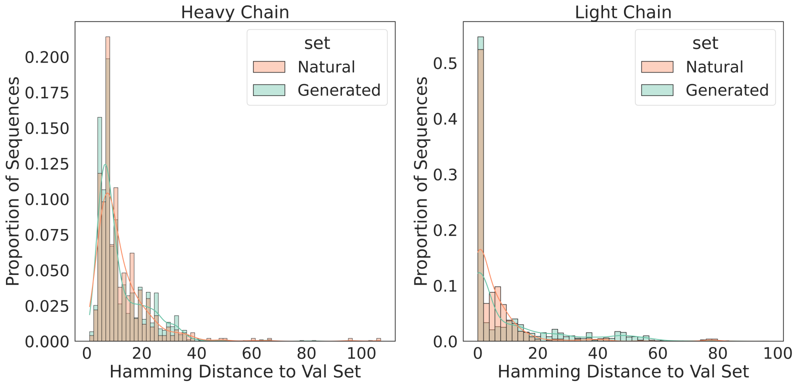


Figure 10: **Sequences generated by developable p-IgGen maintain a similar Hamming distance distribution as seen with p-IgGen.** We calculated the Hamming distance of VH and VL from 2,000 sequences generated by developable p-IgGen with the validation set of developable paired OAS. We also calculated the sequence identity of a random sample of 2,000 developable paired OAS training set sequences to validation set sequences ("Natural").

### 5 Zero-shot Task

For zero-shot prediction, we adapted code from the FLAb repository [Chungyoun et al., 2024] to calculate the perplexity of paired sequences in each dataset and determined the Pearson correlations with the experimental assay data. For IgGen we calculated the mean perplexity of the VH and VL sequences. For the p-IgGen models, we took the perplexity of the concatenated VH and VL sequences.

| Model | Parameters | Pearson Correlation |
|---|---|---|
| ProGen/small | 151M | 0.56 |
| ProGen/medium | 764M | 0.56 |
| ProGen/base | 764M | 0.53 |
| ProGen/xlarge | 6.4B | 0.50 |
| ProGen/large | 2.7B | 0.49 |
| Developable p-IgGen | 17M | 0.42 |
| p-IgGen | 17M | 0.41 |
| IgGen | 17M | 0.28 |
| AntiBerty | 26M | 0.27 |
| IgLM | 13M | 0.27 |
| ProGen/oas | 764M | 0.20 |

Table 2: **The p-IgGen models (p-IgGen and developable p-IgGen) significantly outperform the unpaired IgGen model and other state-of-the-art language models of comparable size for zero-shot expression prediction.** Language models were evaluated for zero-shot prediction of expression levels with a deep mutational scan dataset consisting of 4275 anti-VEGF antibodies [Koenig et al., 2017] using FLAb. Results are ordered by Pearson's correlation (best to worst).

| Model | Parameters | Training Dataset(s) |
|---|---|---|
| AntiBerty | 26M | Unpaired OAS |
| IgGen | 17M | Unpaired OAS |
| p-IgGen | 17M | Unpaired OAS, Paired OAS (finetuning) |
| developable p-IgGen | 17M | Unpaired OAS, Paired OAS (finetuning) |
| IgLM | 13M | Unpaired OAS |
| ProGen/oas | 764M | Unpaired OAS |
| ProGen/small | 151M | UniRef90, BFD30 |
| ProGen/medium | 764M | UniRef90, BFD30 |
| ProGen/base | 764M | UniRef90, BFD30 |
| ProGen/large | 2.7B | UniRef90, BFD30 |
| ProGen/xlarge | 6.4B | UniRef90, BFD30 |
| ESM-IF | 124M | CATH40, UniRef50 |
| MPNN | 1.7M | PDB |

Table 3: **Summary of model parameters and training data.** Inverse folding models (ESM-IF and MPNN) were trained on structural data, while all other models were trained on sequence data. AntiBerty, IgLM, and Progen-OAS were trained on unpaired antibody sequences from OAS [Olsen et al., 2022a]. All other ProGen models were trained on UniRef90 [Suzek et al., 2015], a redundancy-filtered subset of the UniProt dataset, and BFD30, which is mainly from metagenomic sources [Steinegger and Söding, 2018]. ESM-IF was trained on experimental structures from CATH40 [Sillitoe et al., 2015], a redundancy-filtered subset of the Protein Data Bank (PDB) [Berman et al., 2000], as well as AlphaFold2 [Jumper et al., 2021] predicted structures of UniRef40 [Suzek et al., 2015]. MPNN was trained on a subset of experimental structures taken from the PDB.
